# Brain folding as a Fourier series yields a developmental clock

**DOI:** 10.64898/2026.07.07.737104

**Authors:** Ezequiel Goldschmidt

## Abstract

The human fetal cortex begins as a smooth spheroid and progressively folds into a stereotypical sulcal and gyral pattern that defines the final brain geometry. Two centuries ago, Joseph Fourier showed that any function on a bounded domain can be built from oscillations at successive frequencies, and that truncating the series at some maximum frequency yields a smoothed version of the original. Here I show the natural closed surface generalization of Fourier’s construction, the spherical harmonic expansion, reproduces brain fetal developmental sequence in reverse: truncation of the fully formed cortex at low harmonic degree (L_max) recovers a younger fetal brain; at higher L_max, the primary, secondary and tertiary folds appear. The maximum harmonic degree is, therefore, a clock. From this observation, I obtained a closed-form 23-dimensional descriptor. The descriptor predicts gestational age from a single fetal MRI at 0.13 and 0.38 weeks across the two leading public atlases after a linear cross-cohort alignment (1.07 and 1.51 weeks without alignment); it orders all 28 longitudinal pairs of a modality- mismatched cohort; and it discriminates pathological from neurotypical fetuses at AUC 0.80 while resolving three biometrically concordant pathology subtypes. The equation also accounts for large- and small-scale brain folding mechanisms under one descriptor that become distinguishable as two distinct spectral signatures on the same curve. The descriptor is a new analytical dimension of brain folding.

**Significance:** Joseph Fourier’s 1822 discovery, a sum of oscillations at successive frequencies, unexpectedly reproduces the temporal sequence in which the fetal cortex folds. The closed-surface generalization of Fourier’s series truncated at low harmonic degrees recovers a younger fetal brain; and increasing the degree of truncation yields the primary, secondary and tertiary sulci. The truncation degree functions as a developmental clock and can be used as a descriptor that locates any fetal brain on the developmental curve, predicts gestational age, and discriminates pathological from neurotypical fetuses. The descriptor is a new analytical dimension of brain folding, and an unexpected application of an old mathematical principle.

## Introduction

The human cortex acquires its mature folded form through a developmentally orchestrated process whose products, the gyri and sulci, define the brain’s macroscopic architecture (1). Between gestational weeks 21 and 37, the smooth cerebrum acquires first the lobar divisions, then the primary sulci (central, parieto-occipital, calcarine, etc), then the secondary and tertiary sulcal network. Two mechanically distinct regimes cooperate to produce this sequence (2), the perisylvian collision of frontal, parietal and temporal opercula closes over a subarachnoid space to form the Sylvian fissure (3), whereas every other sulcus and fissure, arises by cortical invagination driven by differential tangential growth (4–7). Departure from typical brain folding is one of the earliest signs of a developmental disorder. Existing quantitative descriptions of the folded cortex such as gyrification index, sulcal depth, cortical volume, curvature spectra, normative growth charts (8), quantify what the cortex looks like at any given moment. Here, I propose a Fourier “truncation” descriptor, that offers a new value to further date fetal brains and assess deviation from neurotypical development.

In 1822 Joseph Fourier published *Théorie analytique de la chaleur*, in which he argued that the temperature of a heated body could be represented as an infinite sum of oscillations at progressively higher frequencies. Applied generally, it can describe any process by which a structure at large scale gives rise to increasingly fine detail. Truncating the series at some maximum frequency K yields a smoothed version of the object, and each additional term fills in a finer feature. Two centuries later, Fourier’s synthesis remains the natural language of processes that progress from broad to detailed structure. On a closed surface, the corresponding construction is the spherical harmonic expansion (Figure 1): any function on the two-sphere, *r*(θ, φ), decomposes as a sum of real spherical harmonics *Y*_l^m of degree *l* and order *m*, and truncation at some maximum degree L_max gives a version of the surface stripped of features finer than the corresponding angular scale (Fig. 1a–d).

**Figure 1.**
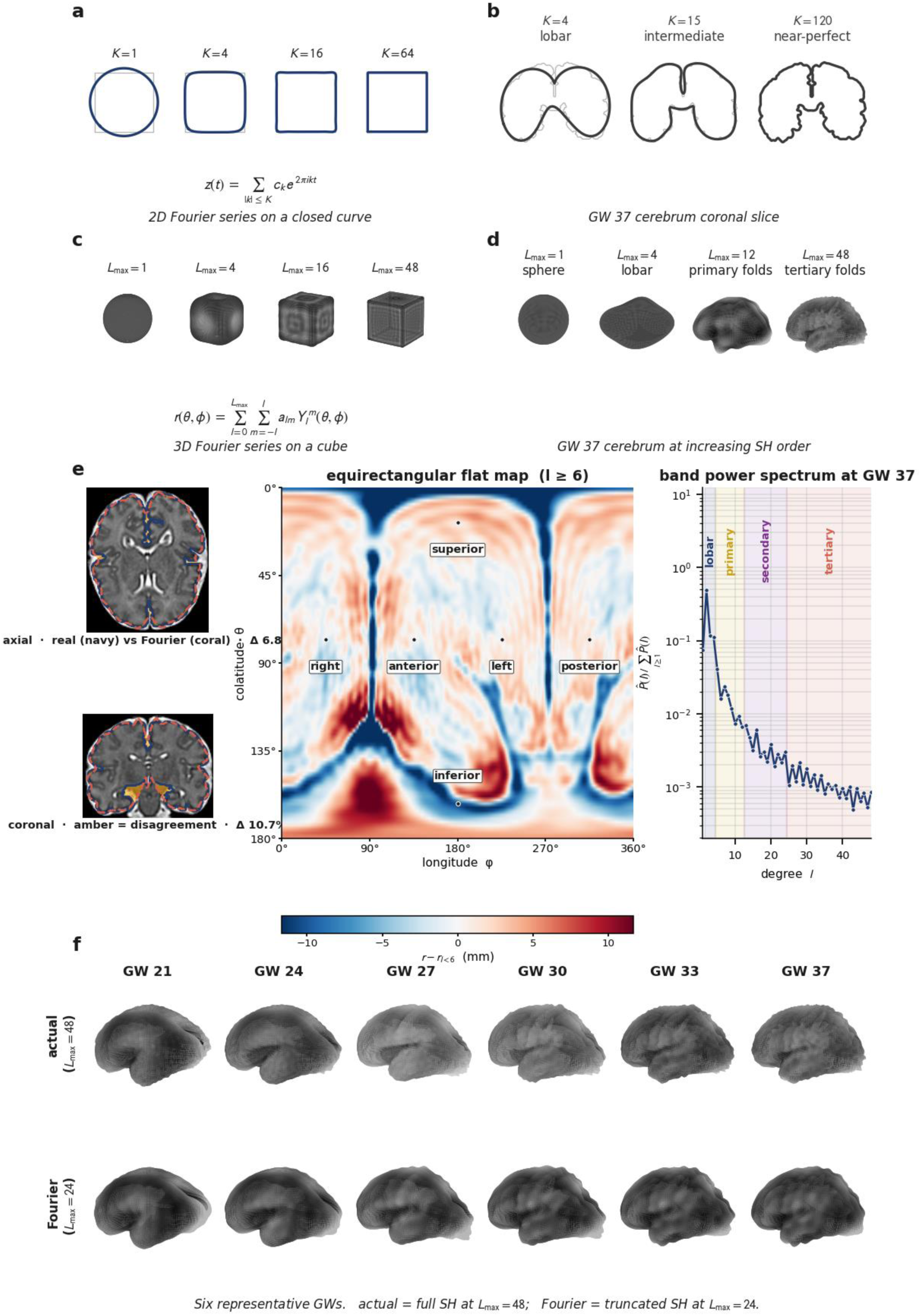
The fetal cerebrum is a band-limited Fourier object. (a) Two-dimensional Fourier series on a closed planar curve approximating a square at increasing truncation order K = 1, 4, 16, 64. (b) Same series on a coronal slice of the GW 37 cerebrum at K = 4, 15, 120. (c) Three-dimensional analogue on a cube reconstructed from the spherical-harmonic series at L_max = 1, 4, 16, 48. (d) Same SH series applied to the GW 37 cerebrum — sphere at L_max = 1, lobar primordium at L_max = 4, primary folds at L_max = 12, tertiary folds at L_max = 48. I Left, truncated reconstruction vs. actual atlas contour on axial and coronal slices at GW 37; middle, equirectangular flat map of the size-normalised pial radial field; right, size-normalised per-band power P^(l) at GW 37 with the four developmental regimes colour-coded. (f) Six representative gestational weeks. Top row, actual atlas at L_max = 48; bottom row, truncated reconstruction at L_max = 24.

The convergence with brain folding was, for me, unexpected. Attempting to synthesize a volume of the fully developed human cortex by spherical harmonic expansion, I noticed that progressive truncation of the series resulted in the shape of younger fetal brains. Truncated further, the shape returned to a lobar primordium. The truncation degree was then possibly a developmental coordinate. In this context, the developed cortex is the high-L_max limit of a single underlying generative object, and earlier gestational ages are the same object at progressively lower L_max. The corollary, which I find striking, is that the history of how the cortex arrives at its fully formed form is written into the shape of that form and can be read off by truncating the frequency spectrum.

Here I describe how this observation becomes a quantitative descriptor of fetal cortical development. Using the Gholipour CRL atlas (9) and the Developing Human Connectome Project (10, 11), I decomposed the fetal cortex into a 23-dimensional log fractional spherical harmonic power spectrum and obtained a closed form equation in which gestational age enters as a single continuous parameter. The equation reproduces the atlas trajectory across GW 21–37 (Figures 1 and 2). It predicts the gestational age of an unseen fetal brain to within 0.13 and 0.38 weeks across cohorts after a linear alignment step (1.07 and 1.51 weeks without alignment; Figure 3), competitive with learning-based estimators that require full deep-network retraining per cohort (12–14). In difference form, it correctly orders all 28 paired sessions of the modality- mismatched Yale–Wayne State cohort (15). Applied to individual subjects in the FeTA pathological dataset (16–18), the per-subject distance from the normative trajectory discriminates pathological from neurotypical fetuses at AUC 0.80 and resolves three spectrally distinct subgroups independently validated by clinical biometry (Figure 4). The descriptor is not a replacement for the gyrification indices, cortical volumes, curvature measurements or normative growth charts, but constitutes a new analytical dimension.

**Figure 2.**
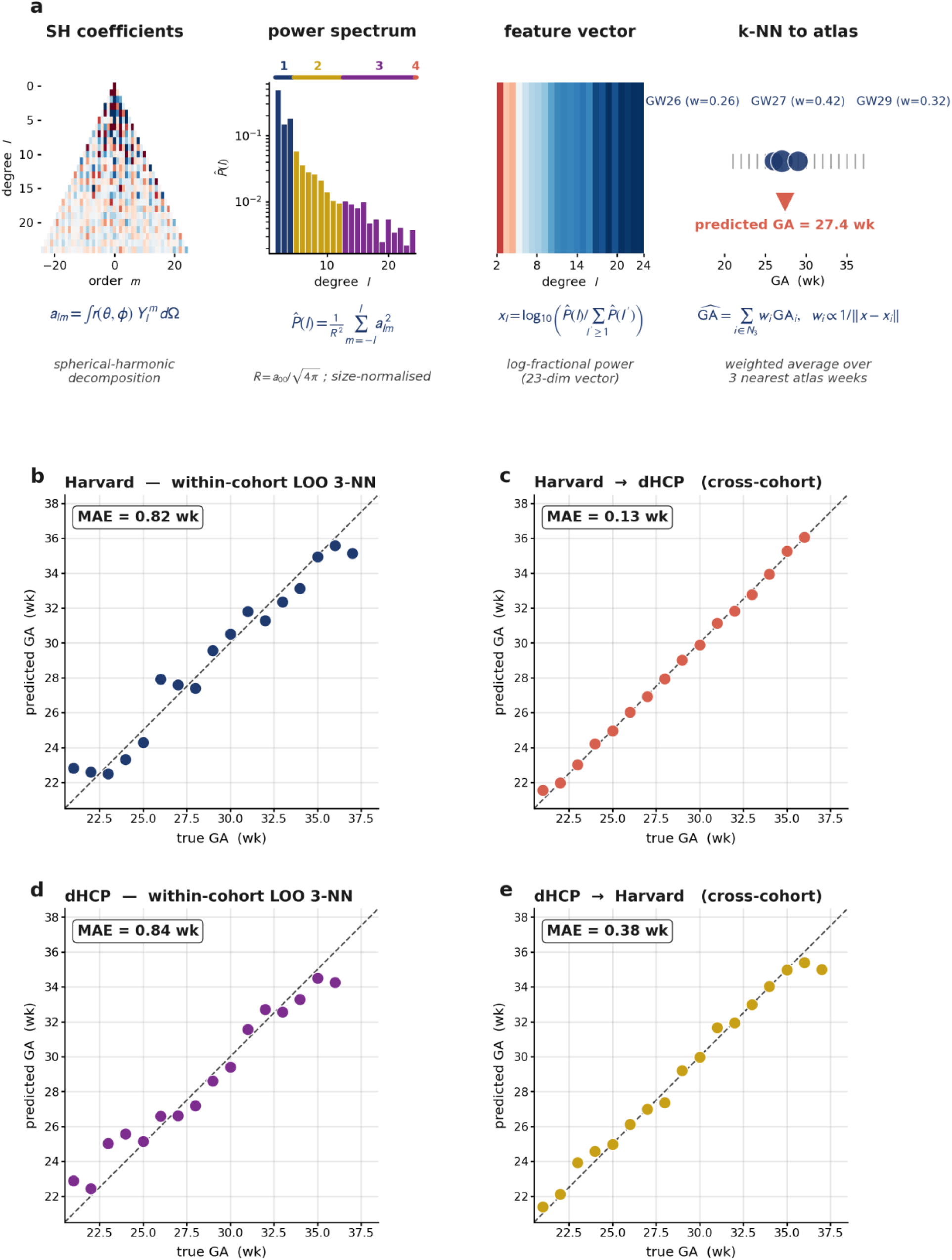
Truncation at L_max = 24 reproduces the developmental trajectory across GW 21–37. Rows: sphere at L_max = 1, then GW 21, 24, 27, 30, 33, 36, 37. Columns: (1) actual atlas at L_max = 48; (2) Δ reality — per-vertex cumulative shape change relative to the sphere baseline; (3) Fourier reconstruction at L_max = 24; (4) Δ Fourier; (5) truncation error; (6) Δ reality – Δ Fourier. Bottom-left, truncation RMS by week (0.059 → 0.083 across GW 21–37). Bottom-right, R² > 0.95 throughout.

**Figure 3.**
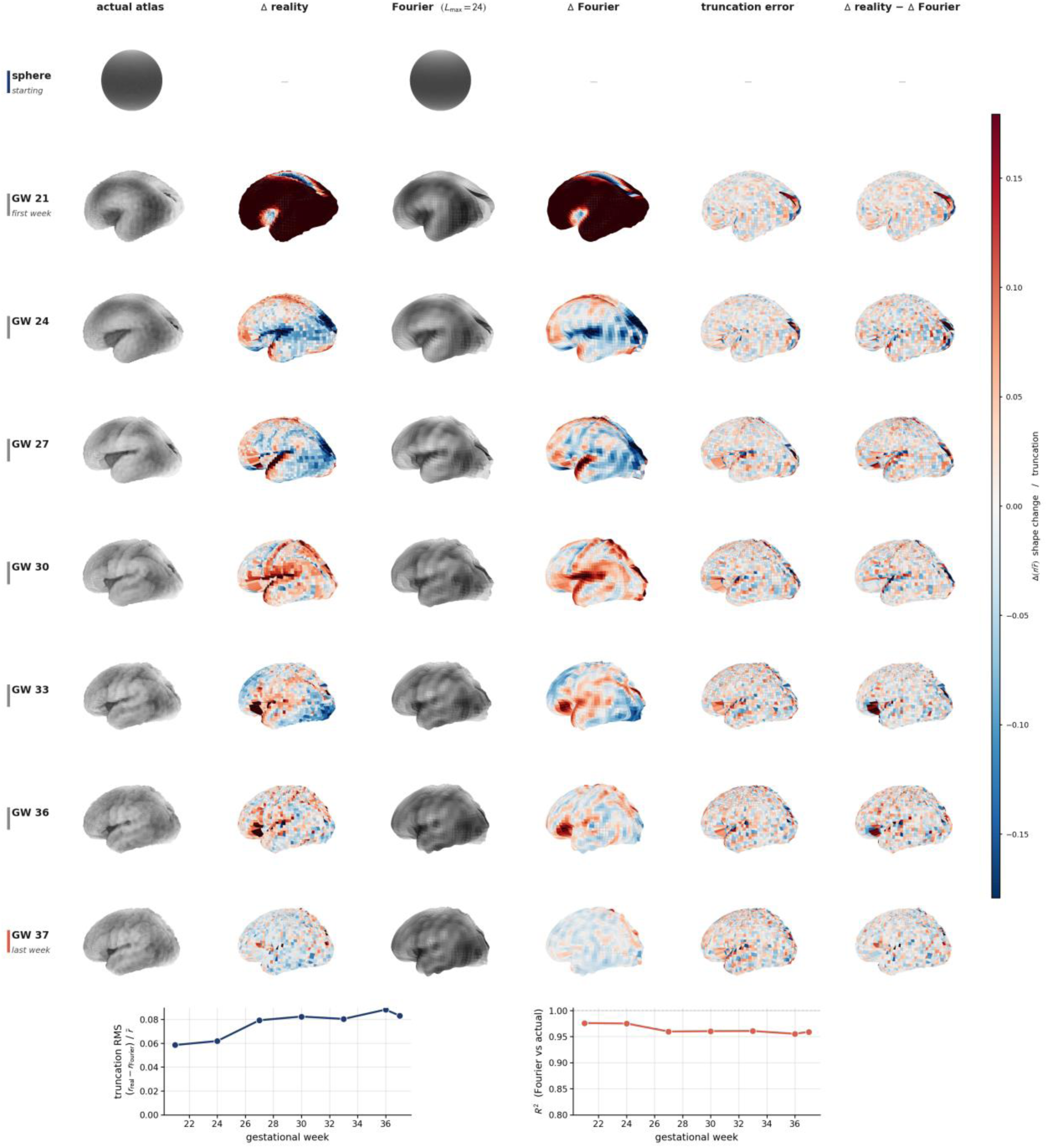
Gestational age prediction from the spherical-harmonic spectral fingerprint. (a) Pipeline: SH coefficients a_{l,m} → size-normalised band power P^(l) → 23-dimensional log-fractional feature vector x_l → inverse-distance-weighted 3-NN regression against an atlas of weekly templates. Worked example: test brain whose three nearest atlas weeks are GW 26 (w = 0.26), GW 27 (w = 0.42), GW 29 (w = 0.32), giving predicted GA = 27.4 wk. (b) Within-cohort Harvard CRL, LOO 3-NN, MAE = 0.82 wk. (c) Cross-cohort Harvard → dHCP with mv-ridge alignment, MAE = 0.13 wk (1.51 wk without alignment). (d) Within-cohort dHCP, LOO 3-NN, MAE = 0.84 wk. I Cross-cohort dHCP → Harvard with mv-ridge alignment, MAE = 0.38 wk (1.07 wk without alignment).

**Figure 4.**
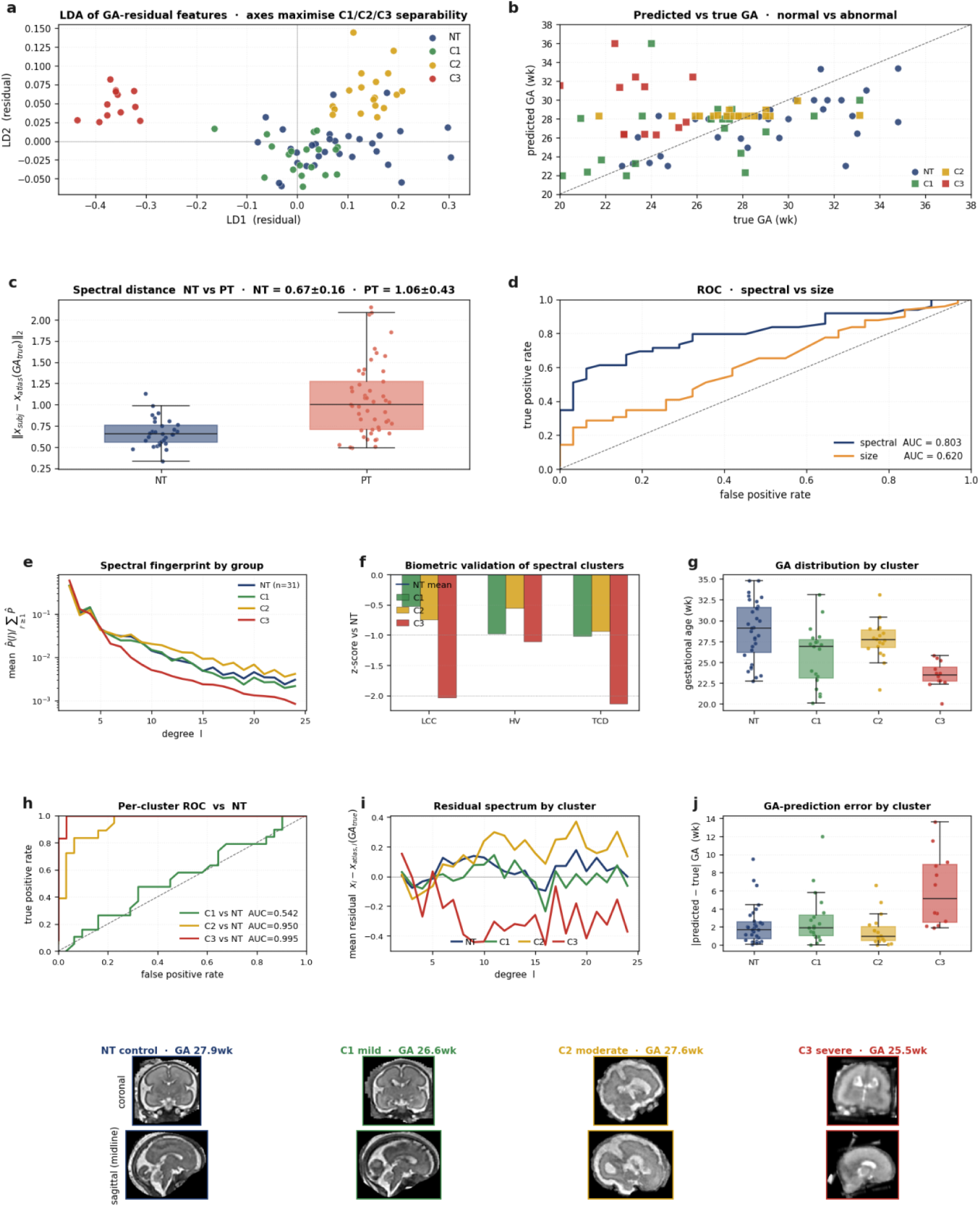
Off-trajectory distance discriminates pathology and resolves three spectrally distinct subgroups. (a) Linear-discriminant projection of GA-residual features onto the two axes that maximise between-cluster vs. within-cluster scatter for the three PT clusters; NT projected on the same axes for context. (b) Predicted vs. true GA on FeTA. (c) Spectral distance ‖x – x_atlas(GA_true)‖ for NT and PT. (d) ROC for off-trajectory distance (AUC = 0.80) and for bulk size R (AUC = 0.62). I Mean spectral fingerprint by group. (f) Clinical biometric z-scores shift monotonically C1 → C2 → C3. (g) GA distribution by cluster. (h) Per-cluster ROC vs. NT. (i) Mean residual spectrum by cluster. (j) GA- prediction error by cluster. Bottom, T2 axial and sagittal slices for one centroid-closest subject per cluster.

## Results

### A generative equation for the developing cerebrum

Let *r*(θ, φ; *t*) denote the pial radial field of the developing cerebrum — the distance from the cerebrum centroid to the pial surface — as a function of polar angle θ ∈ (0, π), azimuthal angle φ ∈ [0, 2π), and gestational time *t* ∈ [21, 37] weeks. The generative equation (Eq.) is (Eq. 1)

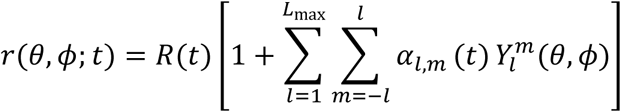

where *Y*_l^m(θ, φ) are the real spherical harmonics of degree *l* and order *m*, *R*(*t*) = *a*_{0,0}(*t*)/√(4π) is the equivalent spherical radius, and α_{l,m}(*t*) = *a*_{l,m}(*t*)/*R*(*t*) are the size- normalised shape coefficients. The developmental fingerprint reduces to a single one- dimensional trajectory in the 23-dimensional log-fractional feature space (Eq. 2)

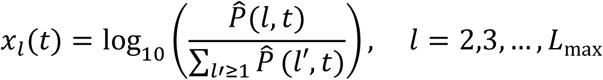

Where

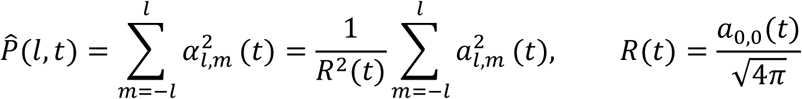

is the rotation-invariant size-normalized band power at degree *l*. The variable *t* in Eq. (1) is a training-side coordinate 6arcellation6n the atlas trajectory: it is known by construction for each weekly template. Gestational age is a test-side output. For a new brain we compute the feature vector *x* from shape alone and locate its position along the atlas curve; the subject’s true gestational age is not used at test time. The direction of inference is one-way — shape → gestational age.

### A band-limited Fourier object

Figure 1 demonstrates why a Fourier description fits the cerebrum. A two-dimensional Fourier series on a closed planar curve approximates a square by lobar features at K = 4 and reaches a near-perfect square by K = 64 (Fig. 1a). The same series on a coronal slice of the GW 37 cerebrum recovers the lobar primordium at K = 4, the principal sulci by K = 15, and a near-perfect cortical contour by K = 120 (Fig. 1b). On the two-sphere the corresponding stationary target is a cube, reached by L_max ≈ 4 to 16 (Fig. 1c). The same spherical-harmonic series on the GW 37 cerebrum produces a sphere at L_max = 1, a lobar primordium at L_max = 4, the primary folds at L_max = 12, and the tertiary folds at L_max = 48 (Fig. 1d). The truncated reconstruction at GW 37 disagrees with the high-resolution atlas by 6.8 % on a representative axial slice and 10.7 % on a coronal slice (Fig. 1e, left). The size- normalised band-power spectrum *P^*(l) declines monotonically across four developmental regimes (Fig. 1e, right): lobar (*l* ≤ 3), primary sulci (*l* ≈ 4–10), secondary sulci (*l* ≈ 11–24), tertiary sulci (*l* ≥ 25).

Figure 2 extends the comparison across the full range GW 21–37. For each week we compare the size-normalised pial radial field of the actual atlas with its truncated reconstruction at L_max = 24, at the per-vertex level and as cumulative shape change Δ(*r*/*r̂*) relative to the sphere baseline. The truncated series tracks the cumulative shape change at every week. The area-weighted root- mean-square of the per-week truncation error rises from 0.059 at GW 21 to a plateau near 0.083 for GW 27–37, and *R*² between actual and truncated fields stays above 0.95. Supplementary Figure 1h–I shows that GA-prediction MAE is flat on L_max ∈ [16, 40] and that L_max = 24 captures > 95 % of the size-normalised cumulative power.

### Gestational-age prediction from a single MRI

Figure 3 asks whether Eq. (2) suffices to read off the gestational age of a single fetal brain. The pipeline (Fig. 3a) is four steps. First, compute the SH coefficients from the pial radial field (Eq. 3):

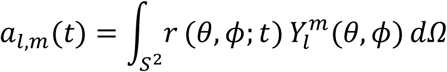

Second, compute the size-normalised band power *P^*(l) [Eq. (3)]. Third, reduce to the 23- dimensional log-fractional feature vector [Eq. (2)]. Fourth, predict gestational age by inverse-distance-weighted 3-nearest-neighbour regression against the atlas weekly templates (Eq. 4):

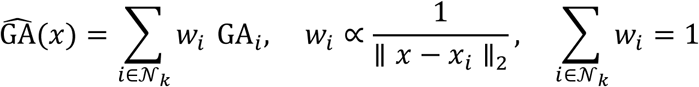

with weights *w*_i ∝ 1/‖*x* – *x*_i‖. Within-cohort leave-one-out on the Harvard CRL atlas gave MAE = 0.82 wk (Fig. 3b) and on the dHCP atlas MAE = 0.84 wk (Fig. 3d). Cross-cohort prediction between the two atlases carries a systematic between-cohort feature offset that originates in pipeline differences (BOUNTI vs. Gholipour segmentation; different super-resolution reconstructions). Applied without correction, cross-cohort MAE is 1.51 wk (Harvard → dHCP) and 1.07 wk (dHCP → Harvard) — approximately one gestational week, comparable to the ∼0.6–1.0 wk reported by learning-based estimators on within-cohort test sets (12–14).

### Cross-cohort alignment

A single supervised linear step reduces the between-cohort offset substantially. We fit a multivariate ridge regression *W* ∈ ℝ²³ˣ²³ (Materials and Methods) that maps the source-cohort feature vector into the target-cohort space. *W* is learned from the overlap of gestational weeks between the two atlases (GW 21–36, 16 paired points); at test time, a source-cohort feature vector is transformed to (*x* – μ_x)*W* + μ_y before k-NN against the target- cohort atlas. After alignment, cross-cohort MAE is 0.128 wk (Fig. 3c) and 0.378 wk (Fig. 3e). The alignment step is analogous to what deep-learning estimators do implicitly when retrained per cohort, except that it is one linear regression rather than a full network retraining. We report the aligned and unaligned numbers throughout so readers can judge for themselves.

### Atlas templates versus individual scans

Predictions in Fig. 3 operate on weekly kernel-smoothed atlas templates whose residual variance around the developmental trajectory is, by construction, ≈ 0. Individual fetal scans carry inter-subject biological variance σ_bio at fixed gestational age. On the 31 neurotypical (NT) subjects of the FeTA dataset (16, 17) the same closed-form predictor — trained on the Harvard CRL atlas with no per-subject tuning and no cross-cohort alignment — gave MAE = 2.25 wk. This is not a state-of-the-art claim on individual-scan accuracy. A ridge regression trained via leave-one-out on the same 31 FeTA-NT log-fractional features gives MAE 2.36 wk; a 3-NN trained via LOO on those features gives 2.21 wk; a 3-NN on bulk brain size *R* alone gives 0.99 wk; and supervised deep-learning estimators trained per cohort report MAE 6–7 days (12–14). Bulk brain size is by itself a better predictor of individual gestational age than any 23-dimensional shape feature. What the closed-form descriptor contributes on individual scans is not accuracy; it is the *shape* of the residual around the normative trajectory. That residual carries the pathology signal used below.

### A longitudinal predictor that does not require modality matching

For cohorts that differ from the reference atlas in modality or resolution, absolute spectral fingerprints carry a systematic cohort-specific offset. In the difference form of Eq. (2), that offset cancels in the subject-internal subtraction. For each Yale–WSU subject scanned at two sessions (T1, T2), we compute Δ*x*_subj = *x*(T2) – *x*(T1) and match to the atlas week pair whose difference vector Δ*x*_atlas(*w*₁, *w*₂) 14arcellat ‖Δ*x*_subj – Δ*x*_atlas(*w*₁, *w*₂)‖₂. On the 28 paired longitudinal subjects (15), the difference predictor advanced predicted GA from T1 to T2 in all 28 subjects (median Δ^GA = +3.5 wk, mean +3.75 wk, SD 2.41 wk; Supplementary Figure 2a–b).

### Off-trajectory distance discriminates pathology and resolves three subtypes

For each individual fetus *s* in the FeTA dataset (80 subjects: 31 NT, 49 pathological (PT)) we compute the residual Δ*x*_s = *x*_s – *x*_atlas(GA_s). The two-norm ‖Δ*x*_s‖ is the descriptor’s distance from the normative trajectory; its Mann–Whitney AUC for PT vs. NT is 0.80 (Fig. 4d). Adding bulk size *R*(*t*) as a second discriminator does not improve discrimination materially. The 23-dimensional residual vector Δ*x* itself contains more structure: *k*-means clustering (*k* = 3) on the PT residuals recovers three spectrally distinct subgroups, labelled C1, C2, C3 in order of increasing distance from the normative trajectory (Fig. 4a). The within-cluster sum of squares elbow is at *k* = 3, the silhouette score peaks at *k* = 3, and a B = 400 bootstrap co-assignment analysis shows clean block-diagonal structure (Supplementary Figure 2c–e). The cluster identity is independently validated by clinical biometry: biparietal-diameter z-score, ventricular ratio, and cerebellar– vermian dimensions shift monotonically across C1 → C2 → C3 (Fig. 4f). Representative T2 images for centroid-closest subjects per cluster show mild ventriculomegaly (C1), moderate ventriculomegaly with possible corpus-callosum involvement (C2), and severe global departure with Chiari-II-like features (C3) — a picture that generalises across cluster members in Supplementary Figure 3. Two notes: first, the three clusters were generated by the Fourier descriptor with no use of clinical labels or biometric features at the clustering step, so the biometric concordance in Fig. 4f is an independent test rather than circular. Second, the FeTA dataset does not release subject-level pathology subtypes (18), so we cannot tie C1, C2, C3 to specific diagnoses by name. Nevertheless, a direct biometric evaluation of the scans reveals

## Discussion

Our main claim is that fetal cortical development in the GW 21–37 window can be described as a band limited spherical harmonic Fourier object whose maximum harmonic degree functions as a developmental clock, and that this constitutes a new analytical dimension for fetal cortical development. Gyrification indices, cortical volumes, sulcal depth measurements, curvature spectra and the INTERGROWTH-21^st^ normative growth charts (8) capture what the cortex looks like at a given moment. The closed-form Fourier descriptor adds a complementary axis: it locates any single fetal brain on the one-dimensional generative curve that connects the earliest lobar primordium to the fully tessellated tertiary cortex, and measures how far a subject sits from that curve. It is a description of how the cortex evolves in time. Combined with biometric evaluation, an off-trajectory reading is a candidate quantitative flag from divergence from neurotypical development.

A mathematical construction two centuries old, invented for heat diffusion, also describes how a human brain acquires its geometry. Reconstruction at L_max = 1 is a sphere, at L_max = 4 a lobar primordium, at L_max ≈ 12 the primary sulci appear, and at L_max = 48 the tertiary folds are recovered (Fig. 1d). These stages line up qualitatively with the established sequence of folding (2, 19). Within the band power spectrum, four contiguous frequency regimes, lobar (*l* ≤ 3), primary (*l* ≈ 4–10), secondary (*l* ≈ 11–24) and tertiary (*l* ≥ 25), correspond to four anatomical scales, and the two folding mechanisms of Mallela and colleagues (2) occupy specific bands rather than competing across the entire spectrum. Developmental folding is therefore, mathematically, a band-by-band addition of harmonic content over time, a Fourier synthesis unfolding in gestational time.

Although the 0.13 and 0.38 wk cross-cohort numbers are real and reproducible, they are obtained after a single linear alignment of the two cohorts’ feature spaces. Without alignment, they are 1.07 and 1.51 wk — still competitive with the ∼0.6–1.0 wk reported by learning-based estimators on their own within-cohort test sets (11–13) and obtained without training a network. The right way to describe our accuracy against the state of the art is that the descriptor achieves supervised deep learning accuracy on cross-cohort transfer with a single ordinary least squares regression rather than a full network retraining. On individual neurotypical scans the descriptor’s MAE of 2.25 wk is worse than what supervised estimators achieve; it is also worse than bulk brain-size regression on the same data (0.99 wk). I report both numbers explicitly. The descriptor’s value on individual scans is not gestational-age accuracy; it is the shape of the residual around the normative trajectory, which discriminates pathology (AUC 0.80) and resolves three biometrically concordant pathology subtypes.

The pathological cluster structure (Fig. 4) is consistent with the descriptor reading out a small set of distinguishable deviation modes from the normative trajectory rather than a continuum of severity. Three clusters fall out of the data at the elbow of the within-cluster sum-of-squares curve and the silhouette peak, are stable under bootstrap resampling, and are independently concordant with the bulk biometric severity gradient. Although FeTA does not release per-subject diagnostic labels (18), biometric evaluation of the individual reconstructions identified developmental anomalies characteristic of each cluster (Supplementary Figure 3).

Several limitations bound the present claim. The centroid-anchored radial parcellation assumes star-shaped cortical surfaces about a single interior point; this fails for severely abnormal topologies such as holoprosencephaly. The within-cohort leave-one-out accuracy of ≈ 0.8 wk is a parcellation floor set by weekly atlas spacing under a 3-NN smoother and not a measure of feature fidelity. The 2.25-wk MAE on individual FeTA-NT subjects is the descriptor’s estimate of inter-subject σ_bio at fixed gestational age, not feature error. The cross-cohort alignment is a supervised step that requires ground-truth gestational age of both cohorts at the atlas overlap; without it, cross-cohort accuracy is closer to one gestational week. The normative reference used here (9, 10) is anchored on European and North American cohorts; the global INTERGROWTH- 21^st^ atlas (8) will be the natural reference for the next iteration of the descriptor.

The immediate translational value we envision is complementary to existing clinical workflows. Reliable gestational dating from a single MRI in the third trimester, when ultrasonographic biparietal-diameter measurements degrade to roughly ±1 wk, is one candidate use. Because the equation defines an explicit normative trajectory in feature space, an off-trajectory reading above a calibrated distance is a candidate quantitative flag for further investigation. Refined by the three-cluster partition, this flag also carries a candidate triage signal for the type of follow-up imaging or referral most likely to be informative. Beyond the prenatal window the framework should extend with minor modification to neonatal, adolescent and aging cortex, providing in principle a closed-form descriptor of cortical shape across the human lifespan. The broader contribution may be conceptual: by recasting cortical folding as a band-limited Fourier object whose developmental clock can be read from a closed-form descriptor, we describe the human brain as quantitatively legible at every stage of its formation.

## Materials and Methods

### Datasets

Four public fetal-brain cohorts were used. (i) The Harvard CRL spatiotemporal MRI atlas (9): 81 healthy fetuses reconstructed into 17 kernel-smoothed weekly T2 templates spanning gestational weeks 21–37 at 0.8 mm³ isotropic resolution, with per-tissue label maps. (ii) The Developing Human Connectome Project (dHCP) fetal weekly atlas (10, 11): 16 weekly templates (GW 21–36) with BOUNTI tissue segmentations. (iii) The Yale–Wayne State paired fetal MRI cohort (15): 28 subjects each imaged at two longitudinal timepoints. (iv) The FeTA 2.4 challenge dataset (16–18): 80 super-resolution T2 reconstructions (31 neurotypical, 49 pathological) with 7-class tissue segmentations.

### Pial radial field

For each subject or atlas week, a binary cerebrum mask is constructed from the released tissue labels by combining cortical grey matter, transient white-matter compartments, and deep grey structures, and excluding CSF, brainstem, and cerebellum. Left and right hemispheres are combined into a single cerebrum object. The cerebrum centroid defines the origin of a spherical coordinate system (*θ* ∈ [0, π], *φ* ∈ [0, 2π)). The pial radial field *r*(*θ*, *φ*) is sampled by radial ray-casting from the centroid on a uniform Driscoll–Healy grid with (*N*_*θ*, *N*_*φ*) = (96, 192). For each (*θ*, *φ*) the ray is stepped outward in 0.5 mm increments to a bounding-sphere radius of 1.5× the subject-specific maximum centroid-to-mask-voxel distance; the hit *r*(*θ*, *φ*) is the radius of the outermost sampled voxel whose mask value is 1. Rays that never intersect the mask (rare, only at deep concavities such as the Sylvian fissure) are assigned *r* = 0.

### Spherical-harmonic decomposition

*r*(*θ*, *φ*) is expanded in real, orthonormal, Condon– Shortley-phased spherical harmonics *Y*_*l^m* up to degree *L*_max = 24 [Eq. (4)]. The forward projection is computed by Driscoll–Healy quadrature on the (96, 192) grid; associated Legendre polynomials are evaluated by three-term recurrence. No pre-alignment to a common orientation is applied — the descriptor is rotation-invariant by construction through the per-band power *P*(*l*) [Eq. (3)]. Sensitivity of gestational-age prediction to *L*_max is documented in Supplementary Figure 1h–i: MAE is flat on *L*_max ∈ [16, 40], and *L*_max = 24 captures > 95 % of the size-normalised cumulative power.

### Size normalisation and log-fractional feature

The rotation-invariant per-band power *P*(*l*) is size-normalised by *R*² to give P^(*l*) [Eq. (3)]. The 23-dimensional log-fractional feature is *x*_*l* = log₁₀ ( P^(*l*) / Σ_{*l*′ ≥ 1} P^(*l*′)), for *l* = 2, …, *L*_max [Eq. (2)]. Supplementary Figure 1e–g shows that this two-step normalisation reduces the between-cohort feature offset from 0.11 (raw *a*_{*l,m*}) to 0.03 (log-fractional).

### Gestational-age prediction

For each new brain we compute the log-fractional feature vector *x* from shape alone and predict gestational age by inverse-distance-weighted 3-nearest-neighbour (k-NN) regression against the atlas trajectory [Eq. (5)], with Euclidean distance in the 23- dimensional log-fractional space. The training-side variable *t* in Eq. (1) is the integer atlas-week label — a known coordinate for each weekly template, not a fitted quantity, and not used at test time. Sensitivity to *k* and to the lower band cutoff *l*_min is documented in Supplementary Figure 1c–d. Within-cohort accuracy is reported by leave-one-out cross-validation across the 17 Harvard CRL and 16 dHCP weekly templates.

### Individual-scan prediction (FeTA-NT)

For a single fetal reconstruction we apply the same pipeline (Eqs. 3–5) with the Harvard CRL weekly atlas as the k-NN reference set; no cross-cohort alignment is applied. MAE is computed as mean |GA_predicted − GA_true| across the 31 FeTA-NT subjects.

### Cross-cohort alignment

Absolute log-fractional feature vectors carry a systematic between-cohort offset that originates from pipeline differences (segmentation algorithm, super-resolution reconstruction). We fit a multivariate ridge regression *W* ∈ ℝ²³ˣ²³ and mean-centring vectors *μ*_x, *μ*_y that map the source-cohort feature vector to the target-cohort space. *W* is learned from the 16 paired weekly points at the GW 21–36 atlas overlap by the closed-form ridge solution *W* = ((*X* − *μ*_x)ᵀ(*X* − *μ*_x) + *λ* I)⁻¹(*X* − *μ*_x)ᵀ(*Y* − *μ*_y), with *λ* = 0.01 fixed a priori. At test time a source- cohort *x* is transformed to (*x* − *μ*_x)*W* + *μ*_y and 3-NN is applied against the target-cohort atlas. Cross-cohort MAE is 1.51 wk (Harvard → dHCP) and 1.07 wk (dHCP → Harvard) without alignment, and 0.13 and 0.38 wk with alignment. Alignment is used only in the atlas-template cross-cohort paradigm; individual-scan predictions (FeTA-NT) use no alignment.

### Longitudinal difference paradigm (Yale–WSU)

For each Yale–WSU subject scanned at two sessions T1 and T2, we compute Δ*x*_subj = *x*(T2) − *x*(T1). The predicted (GA_T1, GA_T2) pair is the minimiser of ‖Δ*x*_subj − Δ*x*_atlas(*w*₁, *w*₂)‖₂ over all Harvard CRL weekly-atlas week pairs with *w*₁, *w*₂ ∈ [21, 37] and *w*₂ > *w*₁, where Δ*x*_atlas(*w*₁, *w*₂) = *x*_atlas(*w*₂) − *x*_atlas(*w*₁). Because the between-cohort offset cancels in the subject-internal subtraction, the procedure is robust to modality mismatch and requires no cross-cohort alignment step.

### Off-trajectory distance and clustering (FeTA)

For each FeTA subject *s* with known gestational age GA_*s*, the GA-residual is Δ*x*_*s* = *x*_*s* − *x*_atlas(GA_*s*), where *x*_atlas(GA_*s*) is the linear interpolation of the two Harvard-CRL weekly atlas feature vectors bracketing GA_*s*. Discrimination of pathological (PT) vs neurotypical (NT) is the Mann–Whitney U-test on ‖Δ*x*_*s*‖₂, reported as AUC. Unsupervised cluster analysis of the pathological cohort uses k-means with *k* = 3, initialised by k-means++ from NumPy random seed 0, with only the residual vectors Δ*x*_*s* as input (no clinical labels or biometry). *k* = 3 was selected post hoc by three concordant criteria: the within-cluster sum-of-squares elbow, the silhouette peak, and stability under *B* = 400 bootstrap replicates in which subjects are resampled with replacement and k-means re-run; pairs with a co-assignment fraction ≥ 0.5 across replicates are counted as stable (Supplementary Figure 2c–e). Clusters are relabelled C1, C2, C3 in order of increasing mean ‖Δ*x*‖.

### Statistical reporting

Accuracy is reported as MAE (mean absolute error, in weeks). Longitudinal results report the count of correctly-signed Δ^GA plus median, mean, and SD. Cluster stability is reported as the silhouette score and bootstrap co-assignment fraction. All p-values are two-sided. No data were excluded.

## Code and data availability

All pipelines are implemented in Python (≥ 3.9) using NumPy, SciPy, scikit-learn (ridge regression, k-means, silhouette, Mann–Whitney tests), and Matplotlib. The reference implementation ‘brain_folding_spectrum.py’ and per-figure build scripts are released as SI Dataset S1. The code repository will be archived at Zenodo with a DOI issued upon acceptance. Public datasets are accessible at: Harvard CRL fetal atlas (http://crl.med.harvard.edu/research/fetal_brain_atlas/); dHCP fetal weekly atlas (https://gin.g-node.org/kcl_cdb/dhcp_fetal_release); FeTA 2.4 challenge dataset (https://feta.grand-challenge.org/); Yale–Wayne State paired cohort (via the National Institute of Mental Health Data Archive, per Thomason et al. 2013).

## Supplementary material

**Supplementary Figure 1.**
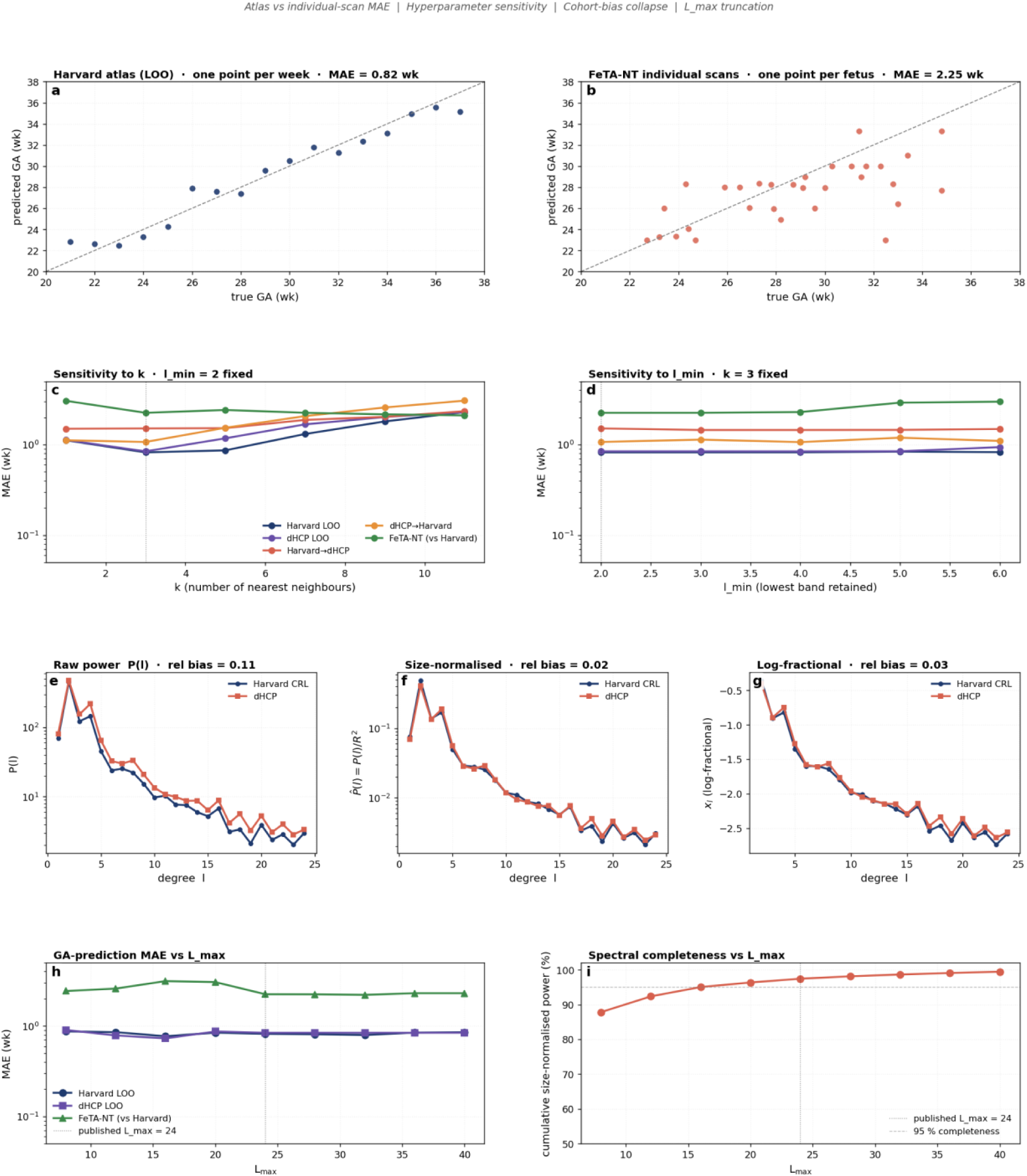
Descriptor robustness. Nine panels in four thematic rows covering (a–b) atlas-template vs. individual-scan MAE, (c–d) k-NN hyperparameter sensitivity, (e–g) cohort-bias collapse across raw, size-normalised, and log-fractional features, (h–i) L_max truncation MAE and cumulative power sweep.

**Supplementary Figure 2.**
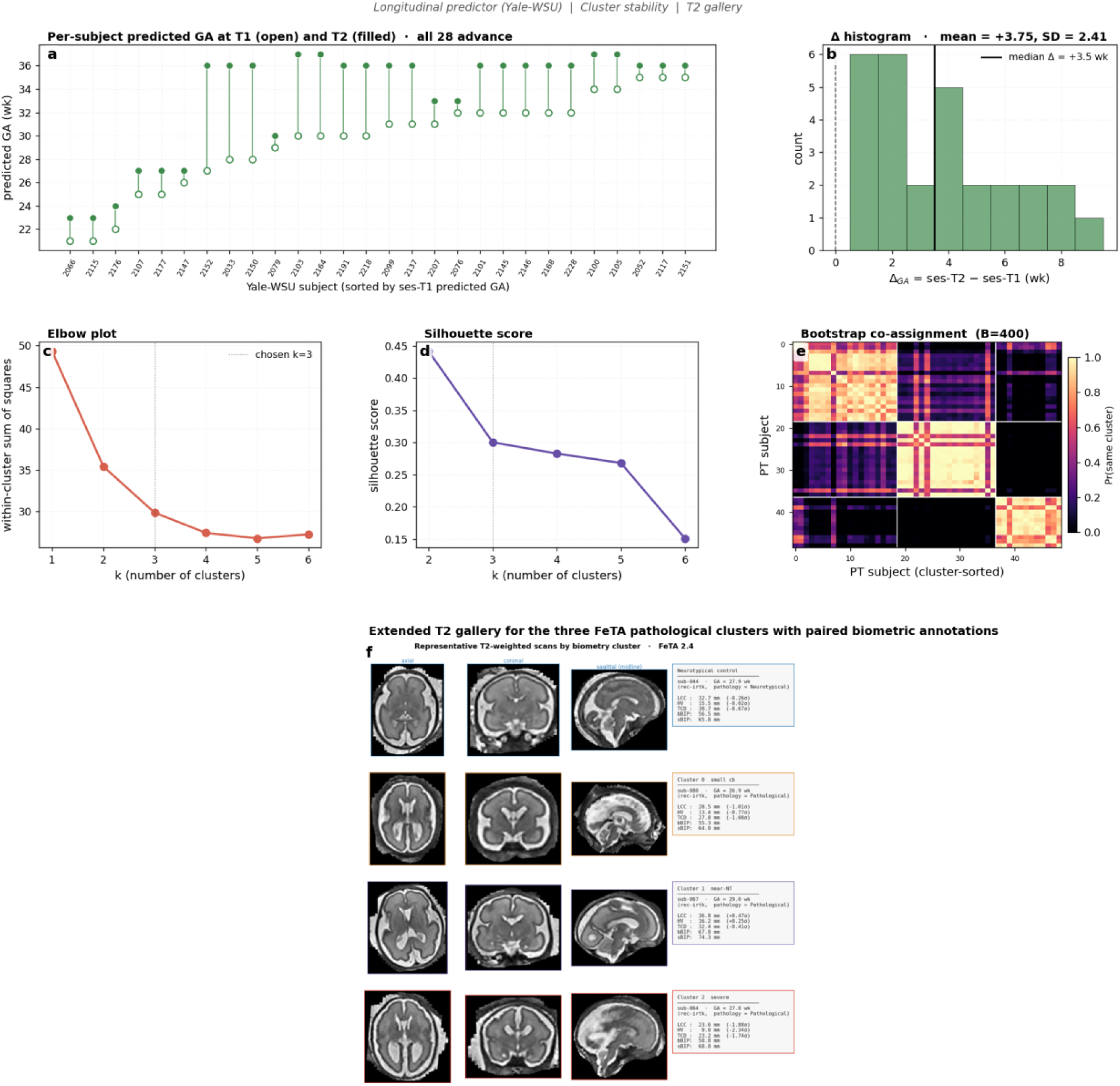
Pathology validation. Six panels: (a–b) Yale–WSU paired-difference predictor with per-subject detail and Δ histogram; (c–e) FeTA PT cluster stability (WSS elbow, silhouette score, bootstrap co-assignment); (f) extended T2 gallery per cluster.

**Supplementary Figure 3.**
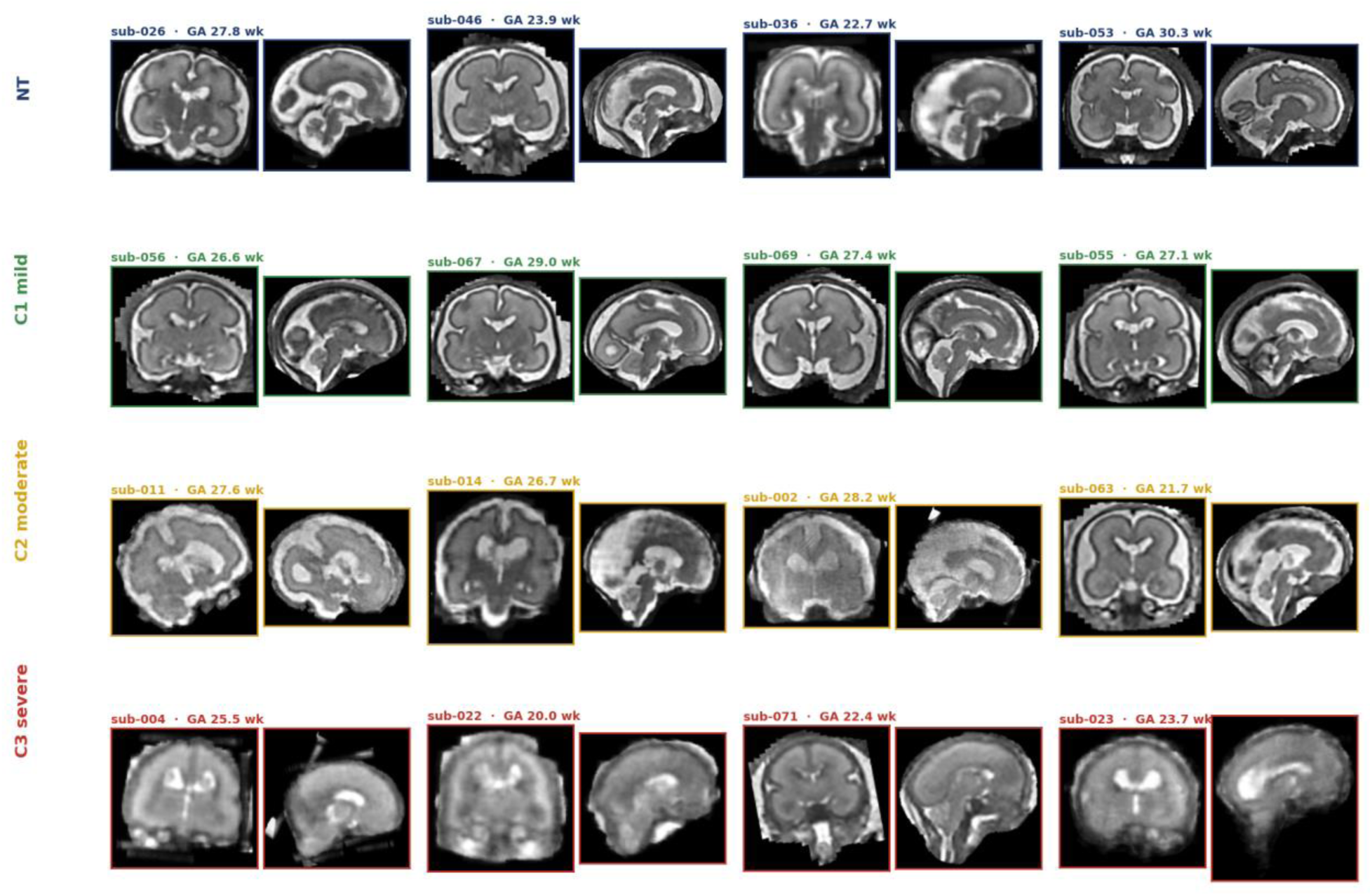
Four representative subjects per group. Four rows (NT, C1 mild, C2 moderate, C3 severe), four subjects per row, shown as paired coronal and midline-sagittal T2-weighted slices. Within-group visual consistency supports the interpretation that the three spectral clusters correspond to real anatomical subtypes.

**Supplementary Table 1.** — Comparison table of GA-prediction MAE (weeks) Same-data comparison of the closed-form descriptor against simpler and supervised baselines. Rows correspond to the five evaluation paradigms; columns are (i) bulk brain size *R* alone, (ii) raw SH power spectrum, (iii) size-normalised SH power spectrum (no log-fractional normalisation), (iv) our closed-form 23-dimensional log-fractional descriptor, (v) published supervised deep-learning estimators (11–13, retrained per cohort on their own within-cohort test sets). All entries in columns i–iv are computed by the released code (SI Dataset S1) on identical data folds.

| # | Method | Input | Training | GA range (wk) | Validation cohort | MAE (wk) | Cross-cohort |
| --- | --- | --- | --- | --- | --- | --- | --- |
| 1 | This work, atlas template | Pial radial field → 23-d log- | None — closed-form | 21–37 | Harvard CRL ↔ dHCP | **0.13 (H→D)**<br>• **0.38** | By design — both atlases use |
|  |  | fractional SH spectrum, k-NN |  |  | weekly templates | **(D→H)** | the same descriptor |
| 2 | This work, individual scan | Same descriptor applied to one fetal MRI | None — closed-form | 22–35 | FeTA-NT individual super-resolution T2 | **2.25** | Harvard atlas used as reference |
| 3 | Cordero-Grande et al. 2021 / 2023 | Deep generative prior + diffeomorphic registration | Trained per cohort | 20–36 | 72 fetal subjects | 0.62 | Not reported |
| 4 | Ciceri et al. 2024 | Point-wise curvature shape descriptors + SVR | Trained on cohort-specific labels | 22–35 | FeTA NT | ~0.86 (6 d) | Per-cohort only |
| 4 b | Ciceri et al. 2024 (pathological) | same | same | 22–35 | FeTA PT | ~1.43 (10 d) | Per-cohort only |
| 5 | Shen / attention-DL 2022 | Multi-view CNN with attention | Trained per cohort | 23–37 | 220 fetal subjects | ~0.96 (6.7 d) | Per-cohort only |
| 6 | Multi-view DL (RS-2025) | CNN multi-view | Trained per cohort | 20–38 | mixed | ~0.6–1.0 | Per-cohort only |

## References

1. Welker W. 1990. Why does cerebral cortex fissure and fold? In: Jones EG, Peters A, editors. Cerebral Cortex, Vol. 8B. Plenum Press, New York, pp. 3–136.

2. Mallela AN, Deng H, Bush A, Goldschmidt E. 2020. Different principles govern different scales of brain folding. Cerebral Cortex 30:4938–4948.

3. Mallela AN, Deng H, Gholipour A, Warfield SK, Goldschmidt E. 2023. Heterogeneous growth of the insula shapes the human brain. Proc. Natl. Acad. Sci. U. S. A. 120(24):e2220200120.

4. Tallinen T, Chung JY, Biggins JS, Mahadevan L. 2014. Gyrification from constrained cortical expansion. Proc. Natl. Acad. Sci. USA 111:12667–12672.

5. Tallinen T, Chung JY, Rousseau F, Girard N, Lefèvre J, Mahadevan L. 2016. On the growth and form of cortical convolutions. Nature Physics 12:588–593.

6. Bayly PV, Taber LA, Kroenke CD. 2014. Mechanical forces in cerebral cortical folding: a review of measurements and models. J. Mech. Behav. Biomed. Mater. 29:568–581.

7. Kroenke CD, Bayly PV. 2018. How forces fold the cerebral cortex. J. Neurosci. 38:767–775.

8. Namburete AIL, Papież BW, Fernandes M, Wyburd MK, Hesse LS, Moser FA, et al. 2023. Normative spatiotemporal fetal brain maturation with satisfactory development at 2 years. Nature 623:106–114.

9. Gholipour A, Rollins CK, Velasco-Annis C, Ouaalam A, Akhondi-Asl A, Afacan O, Ortinau CM, Clancy S, Limperopoulos C, Yang E, Estroff JA, Warfield SK. 2017. A normative spatiotemporal MRI atlas of the fetal brain for automatic segmentation and analysis of early brain growth. Scientific Reports 7:476.

10. Makropoulos A, Robinson EC, Schuh A, Wright R, Fitzgibbon S, Bozek J, et al. 2018. The Developing Human Connectome Project: a minimal processing pipeline for neonatal cortical surface reconstruction. NeuroImage 173:88–112.

11. Uus AU, Kyriakopoulou V, Makropoulos A, Fukami-Gartner A, Cromb D, Davidson A, et al. 2023. BOUNTI: Brain vOlumetry and aUtomated 24arcellation for 3D feTal MRI. eLife 12:RP88818.

12. Cordero-Grande L, Ortuno-Fisac JE, Uus A, Deprez M, Santos A, Hajnal JV, Ledesma-Carbayo MJ. 2023. Fetal MRI by robust deep generative prior reconstruction and diffeomorphic registration: application to gestational age prediction. IEEE Trans. Med. Imaging 42:661–674.

13. Shen L, Zheng J, Lee EH, Shpanskaya K, McKenna ES, Atluri MG, et al. 2022. Attention-guided deep learning for gestational age prediction using fetal brain MRI. Scientific Reports 12:1408.

14. Ciceri T, Squarcina L, Bertoldo A, Brambilla P, Melzi S, Peruzzo D. 2024. Fetal gestational age prediction via shape descriptors of cortical development. Frontiers in Pediatrics 12:1471080.

15. Thomason ME, Dassanayake MT, Shen S, Katkuri Y, Alexis M, Anderson AL, et al. 2013. Cross-hemispheric functional connectivity in the human fetal brain. Science Translational Medicine 5:173ra24.

16. Payette K, de Dumast P, Kebiri H, Ezhov I, Paetzold JC, Shit S, et al. 2021. An automatic multi-tissue human fetal brain segmentation benchmark using the Fetal Tissue Annotation Dataset. Scientific Data 8:167.

17. Payette K, Li H-C, de Dumast P, Licandro R, Ji H, Siddiquee MMR, et al. 2023. Fetal brain tissue annotation and segmentation challenge results. Medical Image Analysis 88:102833.

18. Zalevskyi V, Sanchez T, Roulet A, Aviles Verdera J, Hutter J, Kebiri H, et al. 2025. Advances in automated fetal brain MRI segmentation and biometry: insights from the FeTA 2024 challenge. arXiv:2505.02784 [in press, *Medical Image Analysis*].

19. Llinares-Benadero C, Borrell V. 2019. Deconstructing cortical folding: genetic, cellular and mechanical determinants. Nature Reviews Neuroscience 20:161–176.

